# A meta-analysis of areas of structural variation in grey matter in individuals with Autism Spectrum Disorder (ASD) in relation to gene expression of candidate ASD genes

**DOI:** 10.1101/2021.04.01.438054

**Authors:** Elisa Panzeri, Alessia Camasio, Lorenzo Mancuso, Donato Liloia, Jordi Manuello, Mario Ferraro, Franco Cauda, Tommaso Costa

## Abstract

Autism Spectrum Disorder (ASD) is a set of developmental pathologies with a strong genetic basis and high heritability. Although neuroimaging studies have indicated anatomical changes in grey matter (GM) morphometry, their associations with gene expression remain elusive. In the present study, we aim to understand how gene expression correlates with structural brain aberration in ASD and how it distributes in a functional network perspective. First, we performed an activation likelihood estimation (ALE) meta-analysis to determine GM alteration in the brain, then we selected genes from the SHANK, NRXN, NLGN family and MECP2, which have been implicated with ASD, particularly in regards to altered synaptic transmission. Gene expression maps were built. We then assessed the correlation between the gene expression maps and the GM alteration maps. We found that the default mode network regions were the most significantly correlated with gene expression of selected genes in both areas of GM decrease and increase. The dorsal attention and the cerebellar network regions are significantly correlated with ASD genes. Different networks, namely somatomotor, limbic and basal ganglia/thalamus network - were found in the increase; for each of these networks, however, only a few genes were significant. Our approach allowed to combine the well beaten path of genetic and brain imaging in a novel way, to specifically investigate the relation between gene expression and brain with structural damage, and individuate genes of interest for further investigation in specific functional networks.

## 1. Introduction

Autism spectrum disorder (ASD) is the current diagnostic label to refer to a group of neurodevelopmental conditions affecting approximately 1 in 54 children aged 8 years [1]. This spectrum of disorders is characterized by impairments in social ability and communication, repetitive behaviors, stereotypies and abnormal sensory perception, with varying degrees of severity in phenotype [2].

The etiology of ASD is highly heterogeneous and not fully understood. It is considered to arise from a complex interaction between the genes and the environment [3,4]. According to the current understanding of the field, ASD susceptibility is estimated 40%-80% due to genetic inheritable factors [3,4]. Other responsible factors are the de novo mutation and environmental aspects, for example the age of the mother [3,4]. In the last decade, research in the genetic component of this complex pathology has been fruitful, with hundreds of genes having been identified as causative of ASD [3,4].

ASD has been studied with various approaches, from studies with cellular and molecular resolution, to whole-brain imaging. One such technique is voxel-brain morphometry (VBM), which allows to study brain phenotype in ASD by quantifying morphometric differences between individuals with a certain pathology compared to healthy brain controls [5]. Regions of the brain showing grey matter (GM) abnormalities in volume, density and concentration were identified across the last two decades [6–10]. The genetic bases of ASD have been studied as well. The advent of genome-wide sequencing (GWAS) studies, the process of determining the genomic sequence in its entirety, revealed that the mutations tend to converge in 3 main pathways: synaptogenesis, (i.e. the formation of synapses), WNT signaling (i.e. the process involved with cell fate determination, migration) and neuronal patterning and chromatin remodeling (i.e. the process of changing chromatin architecture to modify access transcription factors’ access to genes) [3,11].

A more comprehensive insight into the disease may be given by combining the genetic and whole brain imaging approach. Studies linking genomic variation to brain fMRI meta-analysis are starting to appear. Grasby et al (2020) [12] combined genetic data and fMRI to link genetic variation to cortical surface area and thickness. They demonstrated that genetic variants associated with brain morphology, are also associated with cognitive function as well as neuropsychiatric diseases [12]. Lau and colleagues (2020) [13] also related gradients of gene expression to brain structure and development. Anomalies at the level of the synapse, thereby leading to altered brain circuitry, have long since been individuated as one of the main underlying mechanisms of ASD [14,15]. Deficit in synaptic transmission could lead to altered connectivity between brain regions [16]. We therefore selected for the present study SHANK3, SHANK1, SHANK2, NLGN3, NGLN4X, MECP2 and CNTNAP2, all of which have a role in synapse formation, maintenance and/or transmission as individuated in a review by [14] as well as in a transcriptomic analysis by [15].

The SHANK family encodes for essential scaffold proteins in the postsynaptic density of excitatory synapses [17], with mutations leading to altered levels of Post-synaptic density (PSD) proteins, synapse morphology and excitatory transmission [17–20]. NLGN3 and NLGN4X encode for post-synaptic cell adhesion proteins, with NLGN3’s mutations having been reported to increase inhibitory transmission and NLGN4X to cause altered excitatory transmission [21–23]. Moreover, defects in NLGN3 and NLGN4X have been reported to lead to impaired synaptogenesis [21]. Pre-synaptic cell adhesion protein NRXN1’s role in synaptic disruption has been attributed to altered Ca2+ entry at the synapse thus impairing neurotransmitter release [14]. Dysfunction in CNTNAP2 has been linked to impaired axonal growth and reduced dendritic arborisation of inhibitory interneurons as well as impaired synaptic transmission [24,25]. MECP2 encodes for a regulator of chromatin remodeling mostly responsible for silencing gene expression, with deficiencies in the gene expression having been reported to induce decreased excitatory transmission due to a decrease in synapse plasticity and number [26,27].

Brain imaging has been a valuable tool since its introduction. However, canonical functional brain imaging relied on recording task-evoked activity. Since then, however, a revolution has occurred, with measurements of intrinsic brain activity being preferred [28]. This intrinsic activity refers to a method that estimates the high-energy interactions between brain regions that occur during the resting state, i.e. when no active task is being performed and therefore the brain is not actively engaged. A number of resting state networks (RSNs), such as the default mode network (DMN), have been thus individuated [28]. Across studies, these networks reveal patterns of activity that are consistent across subjects and reproducible across participants and time [29–32]. Furthermore, they were found to be useful to further our understanding of neurological and neuro-psychiatric disorders [33–37]. In particular, canonical RSNs such as the DMN, salience network (SN), visual and sensorimotor network (SMN) were often found to be functionally or anatomically altered in ASD [38]. Most importantly, RSNs are inheritable, and influenced by genomic factors [39,40]. Thus, RSNs could be a useful tool to interpret the genetic spatial variation associated with ASD morphological alteration.

To our knowledge, no previous research has correlated the genetic expression of genes involved in pathogenesis to areas of anatomical brain damage, as distributed among functional networks. Our approach allowed us to relate gene expression data to anatomical alteration data, to individuate which genes are of interest in which functional network, according to anatomical alteration clusters. Our decision to only select a few representative genes allowed us to explore the results in depth, without dealing with the complexity of big data. Therefore, the present study investigated the spatial distribution of SHANK3, SHANK1, SHANK2, NLGN3, NLGN4X, MECP2 and CNTNAP2 in relation to ASD volumetric modifications and canonical RSNs. The rationale behind our approach was that, being those genes related to ASD pathology, we should expect that their expression in GM areas showing an anatomical alteration would be particularly relevant for the understanding of the disease. Also, we hypothesized that, when assigning those altered regions to canonical RSNs, high-order networks typically associated with ASD such as the DMN would be found to be particularly related to our set of genes [41]. More in general, we expected that the relation between genetic expression of ASD genes and ASD alterations is distributed according to spatial patterns meaningful for brain function and brain functional connectivity.

## 2. Materials and Methods

### 2.1. Selection of gene expression data

The genetic expression data was obtained from the Allen Human Brian Atlas (AHBA). The Allen Human Brain Atlas is one of the projects of the Allen Institute of Brain Science. It is accessible through the Brainmap database (https://human.brain-map.org/). It contains anatomical and histological data - including the RNA microarray data used in this study- collected from six healthy human specimens with no known neurological disease history. 2 specimens contain data from the entire brain, whereas the remaining ones include data from the left hemisphere only. Briefly, the microarray analysis was performed as such: the brains tissues were partitioned into smaller blocks according to their anatomical roles (i.e. if they were cortical or subcortical structures). A minimum amount of tissue was collected from each block and subsequently processed for mRNA isolation. Microarray analysis was carried out by a third-company party (Beckman Coulter Genomics). The data were then normalized to be included in the atlas [42].

Microarray data from the AHBA using the Allen Software Development Kit [43]. We thu*s* obtained gene expression data for specific brain points that we labeled with Talairach coordinates for each brain. For each gene focus we created Voronoi polygons having the gene expression point as the barycenter using the Voronoi tessellation algorithm [44], as done by [45]. Briefly, the Voronoi tessellation is a decomposition of metric space by distances between sets of points. We assigned to each Voronoi polygon the same gene expression value as the barycenter. We thus obtained six brain maps for a single gene- one for each specimen of the AHBA- which we then averaged to obtain a single cerebral map for each gene.

### 2.2. Search and selection of studies

The data for the anatomical study was selected from the BrainMap and Medline databases. The BrainMap database contains a collection of more than 4000 published neuroimaging studies, including both functional and structural (i.e. voxel-based morphometry) metadata [46–49]. The latter was used in this study. The BrainMap database is part of the BrainMap Project and is currently curated by University of Texas Health Science Center San Antonio. Medline is an online database developed by the U.S. National Library of Medicine, containing more that 26 million published works in the field of biomedicine. The data from BrainMap was accessed through the Sleuth 2.4 software package. Two search algorithms were employed as described below to individuate experiments reporting either a voxel-based GM increase or decrease in subjects with ASD:

For GM decreases: *[Experiments Contrast is Gray Matter] AND [Experiments Context is Disease Effects] AND [Subjects Diagnosis is Autism Spectrum Disorders] AND [Experiments Observed Changes is Controls > Patients]*;

For GM increases: *[Experiments Contrast is Gray Matter] AND [Experiments Context is Disease Effects] AND [Subjects Diagnosis is Autism Spectrum Disorders] AND [Experiments Observed Changes is Controls < Patients]*.

To retrieve eventual articles not part of the BrainMap database, we also conducted a systematic search in the Medline database using the MeSH terms as described below:

*(“Autism Spectrum Disorders” [ALL] OR “ASD” [ALL] OR “Autism”) AND (“Voxel-Based Morphometry” [ALL] OR “VBM” [ALL])*.

The systematic search design adhered to the PRISMA Statement international guidelines [50,51]. This study was also compliant with the consensus-based rules for neuroimaging CBMA [52].

We selected peer-reviewed articles containing experiments conducted using whole brain VBM, for which the results were reported either in Taliarach (TAL) or Montreal Neurological Institute (MNI) coordinates. Moreover, the experiments must have reported alteration of GM in ASD patients compared between-group to healthy controls. We then discarded all the experiments which were based either on samples of less than 10 participants, as suggested by [53], or on a ROI analysis, as suggested by [52]. Some studies reported multiple experiments using the same participants. To avoid analyzing the same subjects repeatedly, we selected either the data from the largest experiment or the data that was labeled according to diagnostic subcategories.

Of the initial 517 potential published articles, only 51 fit the inclusion criteria. They included a total of 80 experiments reporting 541 coordinates of GM alterations, subdivided in 244 decrease and 297 increases (see also Figure S1 - PRISMA flow chart). A total of 4849 subjects were included, subdivided in 2366 ASD patients and 2483 healthy controls. The coordinates reported in TAL space were left as they were, whereas we converted all MNI coordinates in TAL space using the ‘icbm2tal’ algorithm [54].

### 2.3. Anatomical likelihood estimation and correlation with gene-expression maps

To evaluate consistent GM alterations across the selected data, we applied the anatomical likelihood estimation (ALE) analysis [55–57], obtaining convergent areas of GM alteration. ALE is a quantitative voxel-based meta-analytical method that allows to estimate consistent areas of activation or morphological alteration across studies [56].

We individuated the brain region presenting values of GM increase and decrease by carrying out two separate analyses. Coordinates were placed in the Talairach space to perform the ALE analysis. The Ginger-ALE software [58] was used to perform the analysis, obtaining two threshold statistical maps of morphological alteration (p < 0.05, min. volume = 150 mm^3^). Each voxel of the map has a value between 0 and 1 showing the probability of an anatomical alteration in the voxel.

We calculated the correlation between ALE maps and each gene expression map, obtaining a correlation map for each ALE-gene pair. We referenced the Yeo extended atlas [59] which subdivides the brain in 7 cortical networks: the Default Mode Network (DMN), the Dorsal Attention Network (DAN), Somatomotor, Frontoparietal, Limbic, Ventral Attention/Salience Network (VAN/SN). Since our results also presented subcortical clusters, we added two other volumes representing the basal ganglia/thalamus (BG/thal) and the cerebellum networks. The BG/thal and cerebellum volumes were obtained from the Brainnetome Atlas [60]. We projected each correlation map in the Atlas and averaged the correlation coefficient of each cluster to obtain its mean alteration-related genetic expression.

We performed a Monte Carlo method to study the significance of our results: we did a random permutation of the correlation clusters (N = 10,000), that is, the value of correlation of each cluster was randomly reassigned to another one. Then, for each iteration, we recalculated each network’s mean correlation value, thus obtaining a distribution of mean correlations for every network. We then calculated the significance threshold with a 95% confidence interval (CI). Genes correlation values falling above the 95% CI were considered to be statistically significant.

## 3. Results

### 3.1. Correlation of decrease clusters

Figure 1 represents the correlation value between gene expression and ALE value for each cluster of GM decrease. As can be seen, decrease clusters are located in several parts of the brain, including the left posterior insula, around left and right central sulcus, along the cingulate cortex, in the bilateral medial temporal cortex, in the precuneus and across the lateral parietal and occipital cortices. The cerebellum shows several clusters of GM decrease as well. Apart, possibly, for a major involvement of posterior regions compared to the prefrontal cortex, the localization of such clusters do not present any meaningful pattern of alteration.

**Figure 1.**
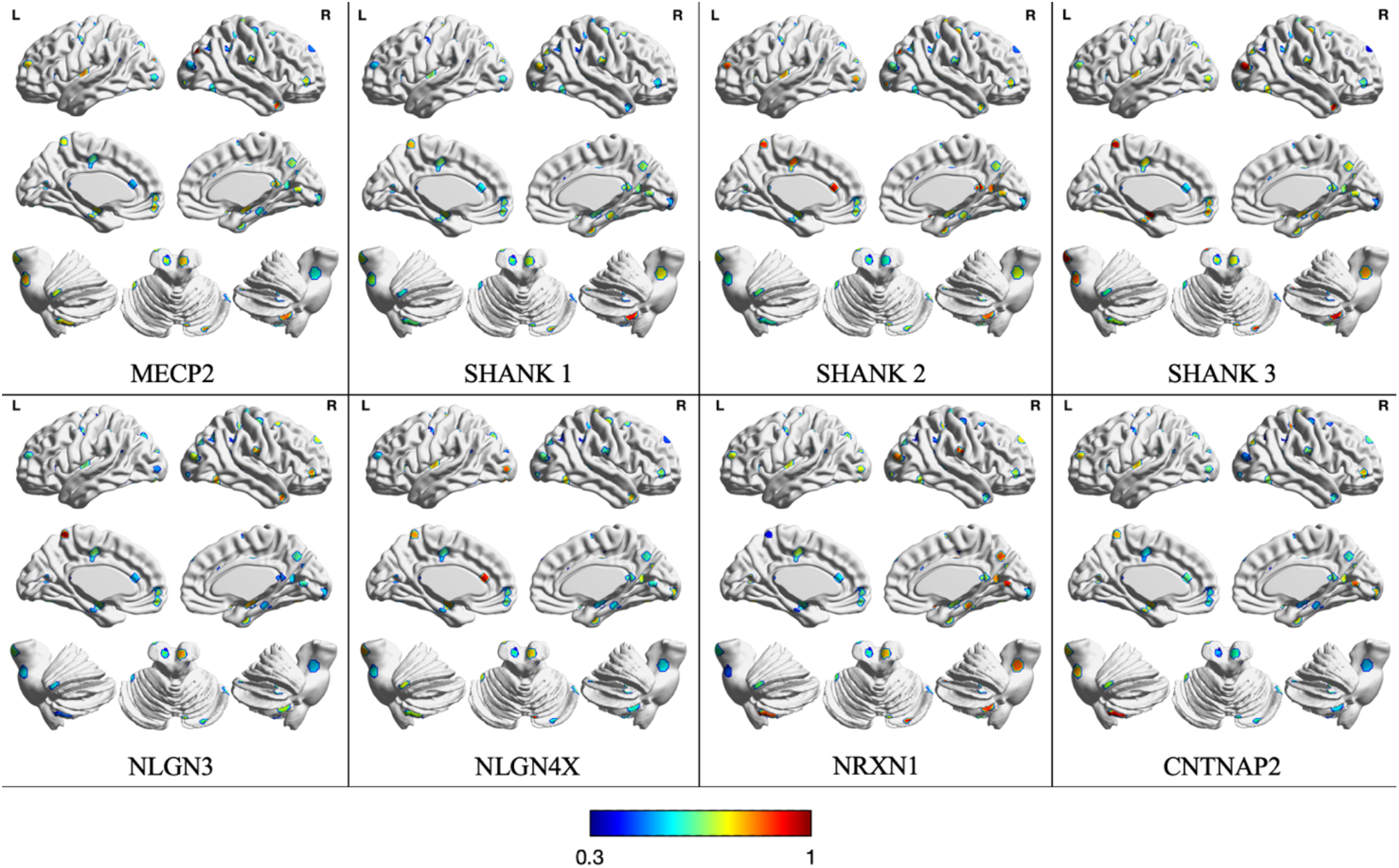
Correlation map between gene expression and ALE-derived GM decrease.

Figure 2 illustrates the average correlation between gene expression and decrease ALE value for each gene and network. The correlation values and significance of each gene for each network are reported in Table 1. Although most genes are expressed homogeneously across networks, the DMN and the DAN make significant exceptions. In the DMN the genes NLGN4X, NRXN1, NLGN3, SHANK1, SHANK3, MECP2 and CNTNAP2 seem to be significantly more correlated than chance (p = 0.05). Similarly, NRXN1, SHANK3 and MECP2 have correlation values significantly higher than the null model in the DAN.

**Table 1.**
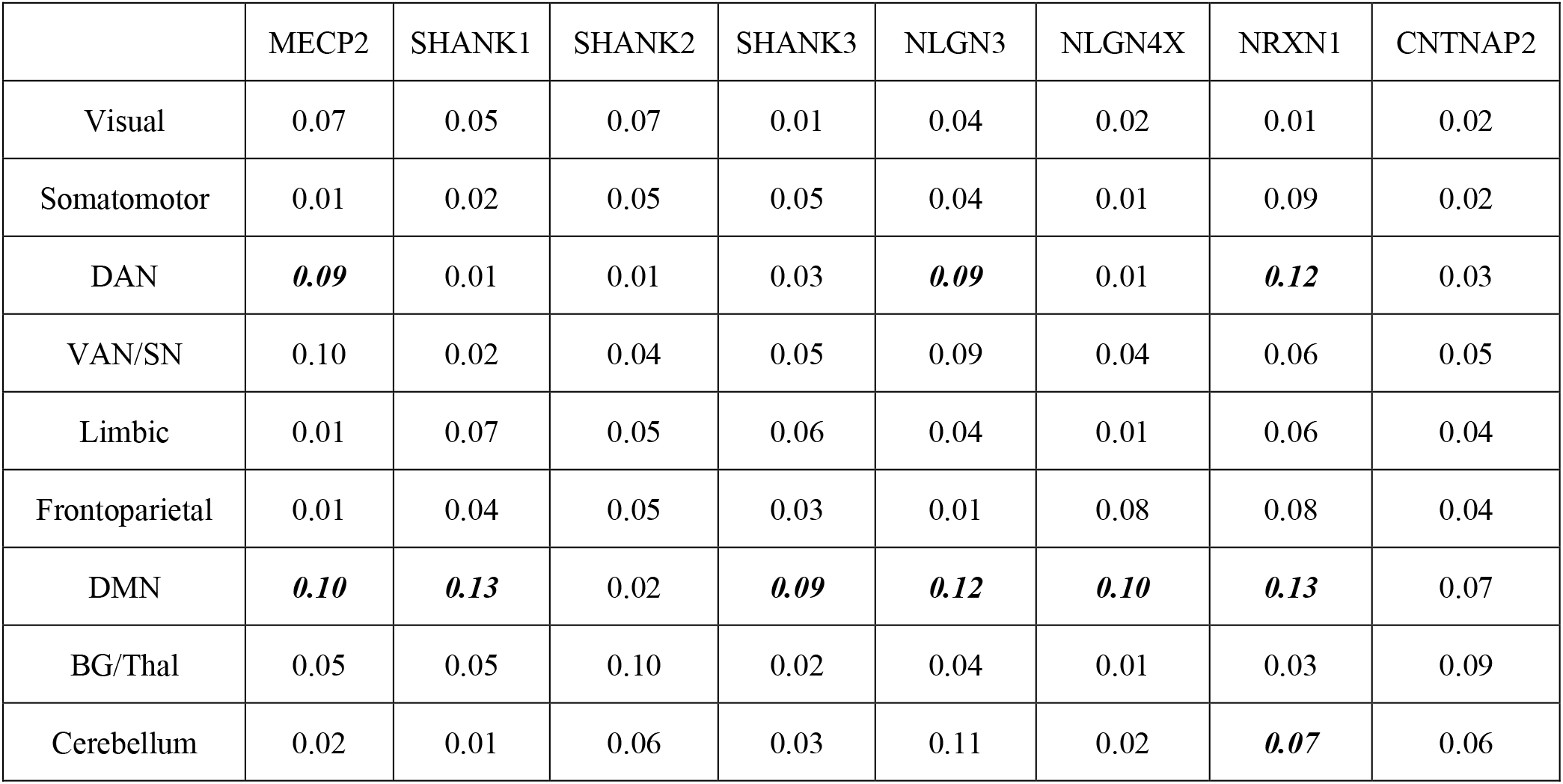
The correlation value of gene expression related to areas of GM decrease in Yeo networks. The numbers in bold indicate significant values. DAN: Dorsal Attention Network, VAN/SN: Ventral Attention/Salience Network, DMN: Default Mode Network, BG/Thal: Basal Ganglia/Thalamus

|  | MECP2 | SHANK1 | SHANK2 | SHANK3 | NLGN3 | NLGN4X | NRXN1 | CNTNAP2 |
| --- | --- | --- | --- | --- | --- | --- | --- | --- |
| Visual | 0.07 | 0.05 | 0.07 | 0.01 | 0.04 | 0.02 | 0.01 | 0.02 |
| Somatomotor | 0.01 | 0.02 | 0.05 | 0.05 | 0.04 | 0.01 | 0.09 | 0.02 |
| DAN | **0.09** | 0.01 | 0.01 | 0.03 | **0.09** | 0.01 | **0.12** | 0.03 |
| VAN/SN | 0.10 | 0.02 | 0.04 | 0.05 | 0.09 | 0.04 | 0.06 | 0.05 |
| Limbic | 0.01 | 0.07 | 0.05 | 0.06 | 0.04 | 0.01 | 0.06 | 0.04 |
| Frontoparietal | 0.01 | 0.04 | 0.05 | 0.03 | 0.01 | 0.08 | 0.08 | 0.04 |
| DMN | **0.10** | **0.13** | 0.02 | **0.09** | **0.12** | **0.10** | **0.13** | 0.07 |
| BG/Thal | 0.05 | 0.05 | 0.10 | 0.02 | 0.04 | 0.01 | 0.03 | 0.09 |
| Cerebellum | 0.02 | 0.01 | 0.06 | 0.03 | 0.11 | 0.02 | **0.07** | 0.06 |

**Figure 2.**
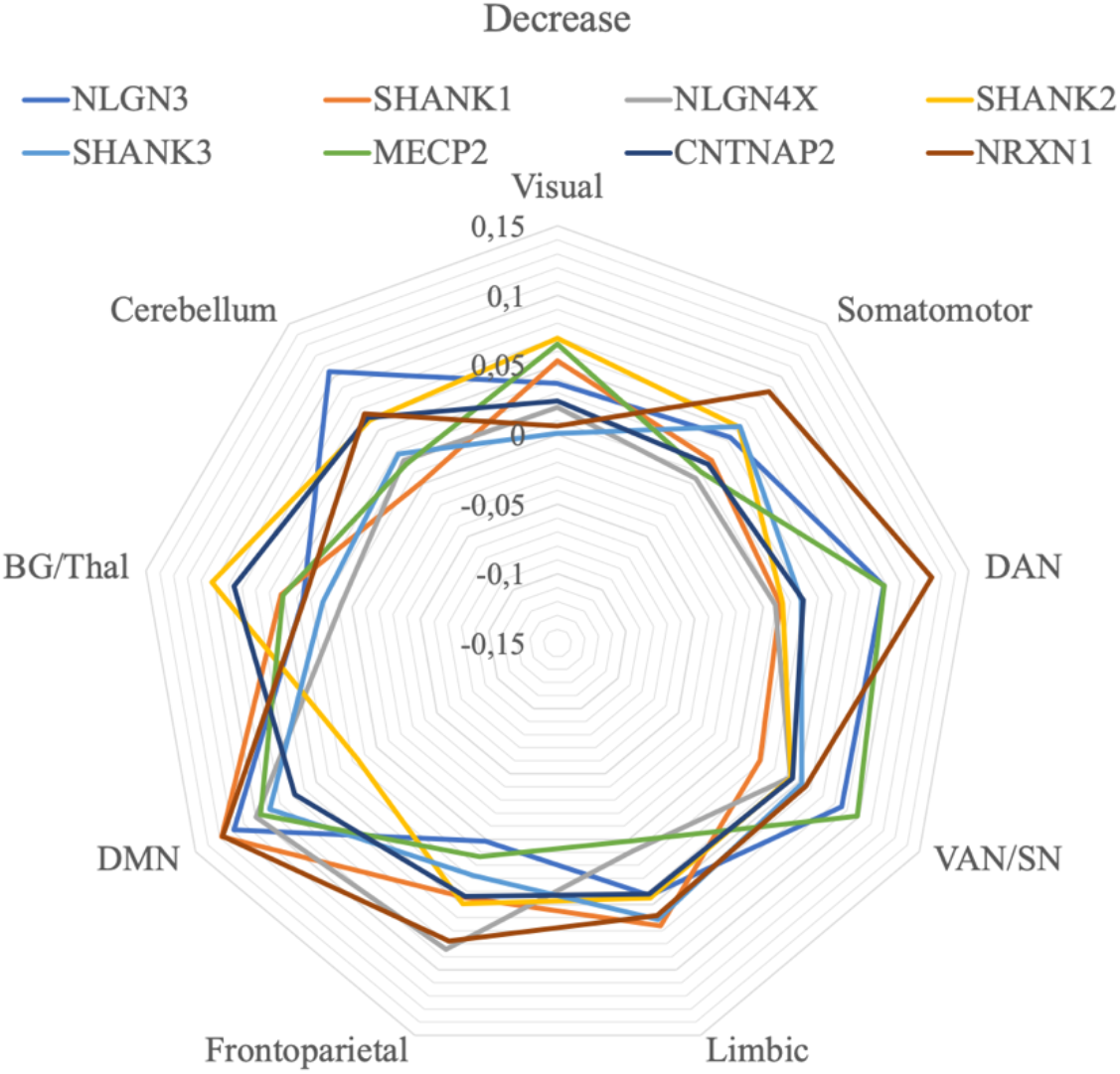
The correlation between gene expression values and GM decrease in the Yeo networks. Both significant and non-significant values are represented. DAN: Dorsal Attention Network, VAN/SN: Ventral Attention/Salience Network, DMN: Default Mode Network, BG/Thal: Basal Ganglia/Thalamus

Also, the cerebellum has one significantly correlated gene (NRXN1, r = 0.07).

### 3.2. Correlation of increase clusters

The correlation values between the gene expression and the GM increase ALE values are represented in Figure 3, while the average correlations of each network for each gene are shown in Figure 4. Here, the lateral prefrontal and temporal cortices are more involved than in the decrease maps, while the parietal and occipital lobes are relatively preserved. Compared to the decrease maps, the cerebellum seems to be less altered as well. Curiously, the midline cortical structures of the left hemisphere show several clusters of increases, while their homotopic counterparts display less alterations of the same kind. Apart from these observations, the various genes show different local correlations, with no obvious pattern to report.

**Figure 3.**
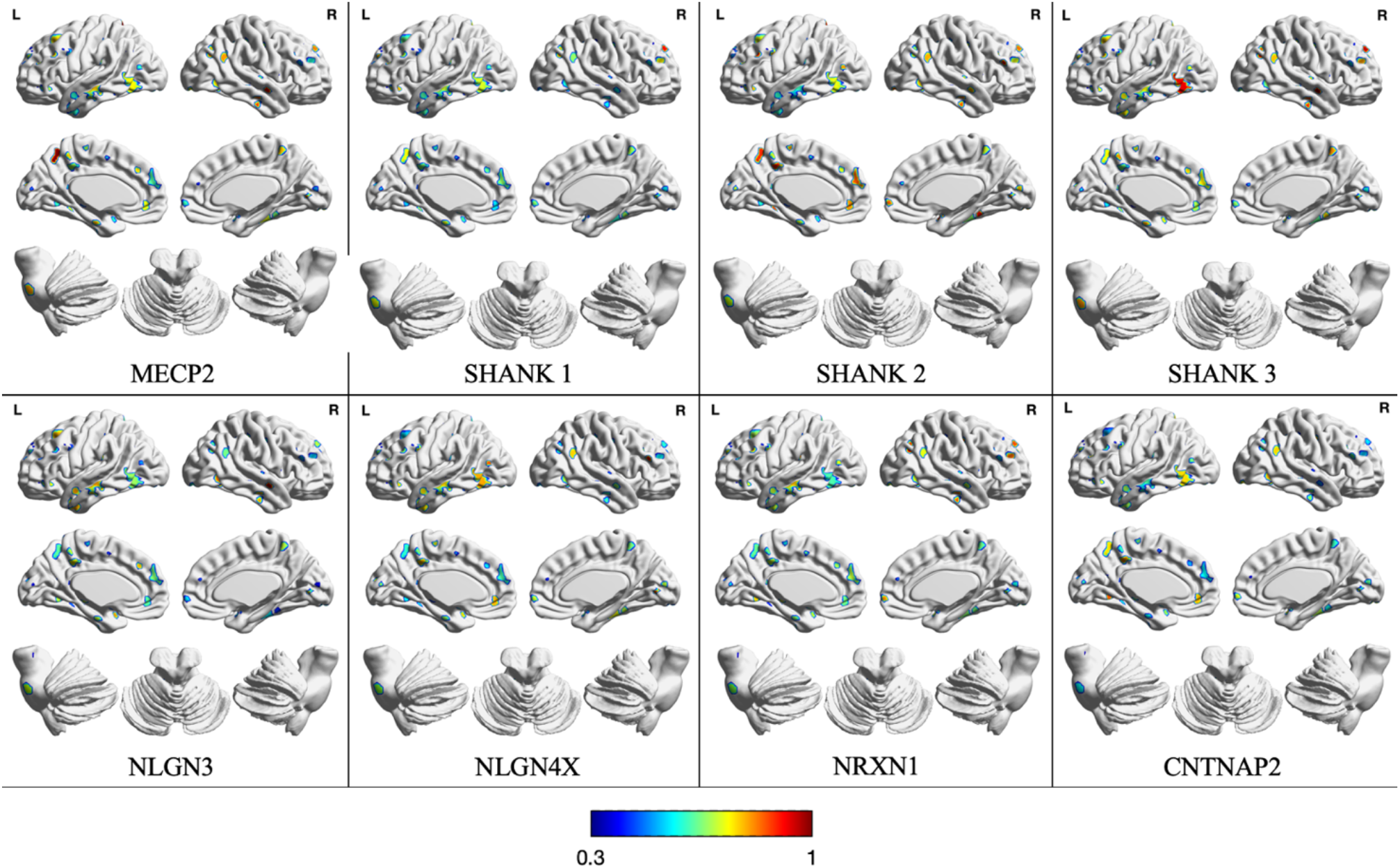
Correlation map between gene expression and ALE GM increase.

**Figure 4.**
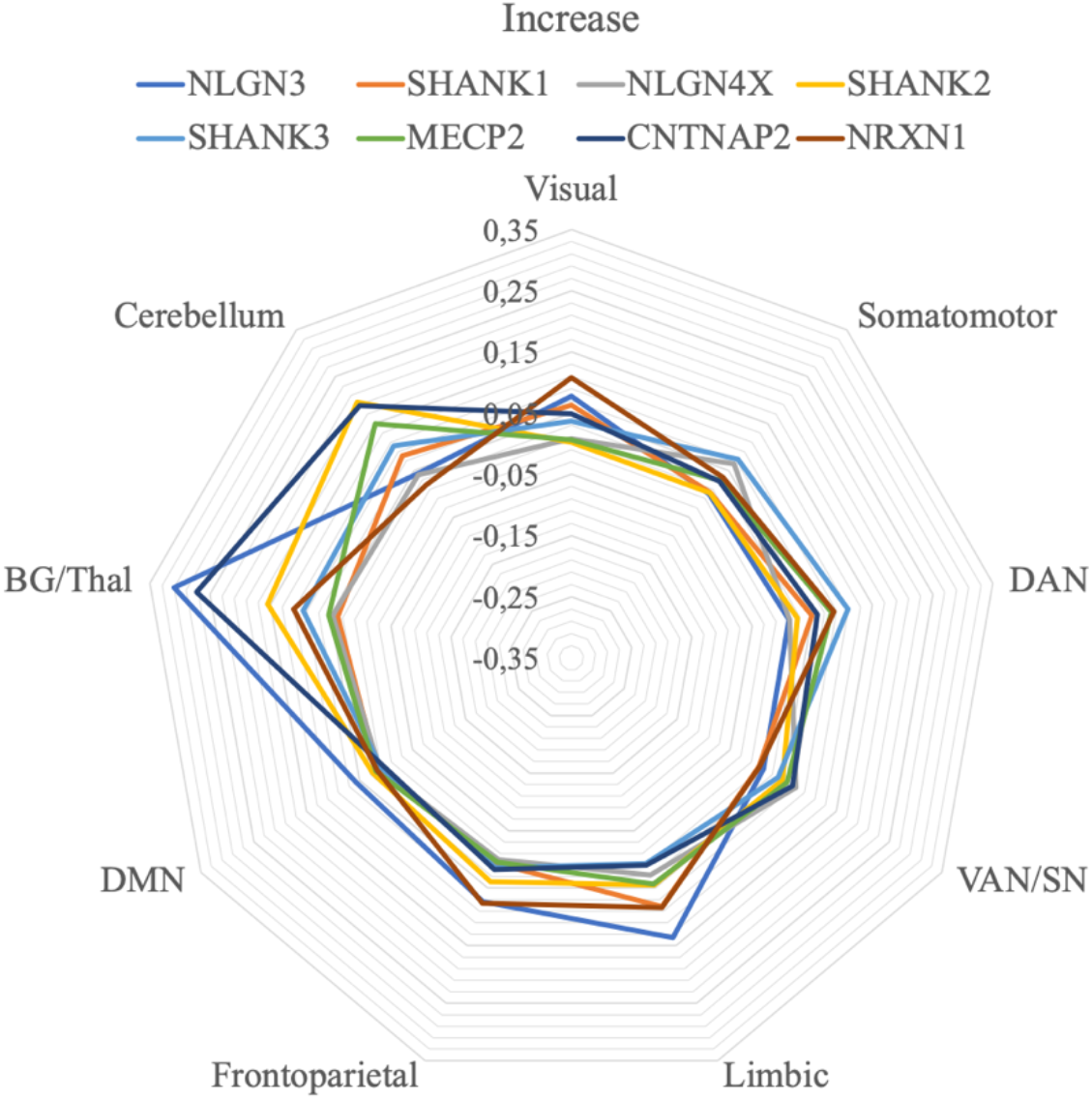
The correlation between gene expression values and GM increase in the Yeo networks. Both significant and non-significant values are represented. DAN: Dorsal Attention Network, VAN/SN: Ventral Attention/Salience Network, DMN: Default Mode Network, BG/Thal: Basal Ganglia/Thalamus

As can be seen in the radar plot in Figure 4, most genes correlate similarly with the increase ALE values of the various networks, with the exception of the BG/THAL and the cerebellum. However, such a visible trend is not confirmed by the Monte Carlo procedure. The significance of the correlation values are represented in Table 2. Compared to the decreased results, there are less significant values per network. The cerebellum is not significantly more correlated to any gene than chance. The discrepancy between the Monte Carlo test and the average cerebellar values observed in the radar graph is likely due to the fact that such structure is characterized by only two increase clusters. The networks revealing at least an average correlation between gene expression and GM increases are the somatomotor (NLGN4X, r = 0.06), the limbic (NLGN3, r = 0.14), the DMN (NLGN3, r = 0.06) and BG/Thal (NLGN3, r = 0.31; CNTNAP2, r = 0.27). In short, NLGN3 is the only gene significantly correlated with increases in more than one network, and BG/Thal is the only system presenting more than one significant correlation. In any case, it might be worth noting that the DMN is the only functional system showing significant values in both states of GM alteration (decrease and increase), highlighting a potentially significant role of the network in the pathological structural alterations found in ASD.

**Table 2.** The correlation value of gene expression related to areas of GM increase in Yeo networks. The numbers in bold indicate significant values. DAN: Dorsal Attention Network, VAN/SN: Ventral Attention/Salience Network, DMN: Default Mode Network, BG/Thal: Basal Ganglia/Thalamus

|  | MECP2 | SHANK1 | SHANK2 | SHANK3 | NLGN3 | NLGN4X | NRXN1 | CNTNAP2 |
| --- | --- | --- | --- | --- | --- | --- | --- | --- |
| Visual | 0.01 | 0.06 | 0.01 | 0.04 | 0.08 | 0.01 | 0.11 | 0.05 |
| Somatomotor | 0.03 | 0.01 | 0.01 | 0.07 | 0.01 | **0.06** | 0.04 | 0.03 |
| DAN | 0.08 | 0.05 | 0.03 | 0.11 | 0.01 | 0.01 | 0.09 | 0.06 |
| VAN/SN | 0.06 | 0.01 | 0.05 | 0.04 | 0.01 | 0.07 | 0.01 | 0.07 |
| Limbic | 0.04 | 0.08 | 0.05 | 0.01 | **0.14** | 0.03 | 0.08 | 0.01 |
| Frontoparietal | 0.01 | 0.01 | 0.04 | 0.01 | 0.07 | 0.01 | 0.08 | 0.02 |
| DMN | 0.02 | 0.02 | 0.03 | 0.01 | **0.06** | 0.01 | 0.02 | 0.01 |
| BG/Thal | 0.05 | 0.04 | 0.15 | 0.10 | **0.31** | 0.05 | 0.11 | **0.27** |
| Cerebellum | 0.15 | 0.08 | 0.20 | 0.10 | 0.04 | 0.04 | 0.02 | 0.19 |

### 4. Discussion

In the present research, we studied the correlation between genetic and structural data in various RSNs. First, we conducted a coordinate-based meta-analysis study to locate the areas of structural alteration common to ASD patients with the ALE methodology. Then, we correlated gene expression data in the individuated regions in various brain networks and tested their significance with a Monte Carlo method. Our main finding was that the DMN was significantly correlated with gene expression of at least one candidate ASD gene in both decrease and increase analyses. Another network that deserves mention because of its significant correlation in the decrease maps was the DAN, while in the increase only the BG/Thal might be considered somewhat meaningful.

Regarding GM decrease results, as already mentioned, the DMN correlates significantly with six out of eight of the genes investigated in the present study. The DMN is a resting state brain network that is associated with social cognition, episodic memory and Theory of Mind [61]. Dysfunction in the connectivity of the DMN has been linked with ASD, especially in regards to dysfunction of social information processing, particularly in regard to oneself [41,62]. Moreover it is linked with an inability to read mental states of others [41,62]. Our results confirm and extend previous literature, as they highlight the DMN as a network of particular interest in the pathology of ASD. Moreover it identifies MECP2, SHANK1, SHANK3, NLGN3, NLGN4X and NRXN1 as significantly expressed, therefore opening up a path of investigation into how altering their function and expression in specific areas that are functionally connected in the DMN may be reflected in pathogenic anatomical changes.

We also found that the DAN is significantly correlated with three genes in the decrease but not increase data. The DAN is of particular interest in the pathology in ASD as it deals with attention, deficits of which are one of the most consistent symptoms across individuals in the ASD spectrum [63]. Our results supplement a study by [64]. They found that adults with ASD display lower functional connectivity between nodes of the DAN compared to neurotypical controls. Interestingly, we found significance for such networks only in the decrease analysis, highlighting a possible correspondence between hypo-connectivity GM morphometrical reductions.

We also found that in the decrease there was a significant correlation in the cerebellum, in accordance to the majority of studies, which shows a hypo-connectivity between the cerebellum and the cortical regions in individuals with ASD [65].

With respect to GM increase, we found a minor number of significant correlations between genes and networks compared to the decrease analysis. This could be linked to the fact that GM increase patterns show a faster temporal development than the GM decrease [34]. In any case, it might worth mentioned that the somatomotor, limbic, DMN and BG/Thal presented at least a significant correlation, in accordance with studies reporting an over-connectivity in each one of these networks [66,67], again supporting a correspondence between the direction of connectivity changes and volumetric modifications.

As regards the genes investigated, NLGN3 was the gene with the most overall significant correlations when considering both decrease and increase together ASD cognitive deficits have been hypothesized to be correlated with altered NLGN family function [14,15]. More specifically, NLGN3 encodes for a cell adhesion protein that is fundamental for proper synaptic transmission. The relation between this gene and ASD is supported by electrophysiological and molecular studies. For example, a study by [68] found that a common ASD-mutation in NLGN3 decreases synchronicity between brain regions and affects axon architecture complexity, thereby decreasing neuronal connectivity [68].

NLGN4X, which we found to be significantly associated to DMN in the decreases and to the somatomotor network in the increases analyses, is known to be involved with ASD phenotypes by altering excitatory synaptic transmission due to its location in the postsynaptic density [22,69]. However, it’s role in long range connectivity and transmission of electric signaling is still not well understood, in part owing to the fact that NLGN4-like protein in mice co-localizes at inhibitory synapses, instead of excitatory as in humans [69]. Further insight into the role of this protein in the transmission of electric signaling would be beneficial in understanding the correlation with the areas of GM alteration found in the present study.

The neurexin superfamily (consisting, in our data, of NRXN1 and CNTNAP2) was also found to be significantly correlated with areas of altered GM in the DAN, DMN, Cerebellum networks and the BG/Thal networks. CNTNAP2 is found mostly at the nodes of Ranvier and is involved with the electrochemical transmission in neurons [70]. Various CNTNAP2 polymorphisms have been found to predispose to ASD via altered long-range connectivity in the brain [70]. NRXN1 is a presynaptic protein which forms complexes with various post-synaptic proteins including the neuroligin family. NRXN1 heterozygous mutations have been found to reduce the brain metabolism and thus reduce the efficiency of connectivity in neural networks including thalamic, mesolimbic and cortical systems [71]. As previously mentioned, long-range disconnections between brain regions are often thought to be the etiology of neurodevelopmental disorder including autism[72–75].

We found that the SHANK family of scaffolding post-synaptic proteins to be significantly correlated to some canonical networks only in the decrease analysis. SHANK3 ASD-causative loss of functions mutations in particular has been found to be associated with low functioning autism, with both intellectual and language disability, on top of the social deficit [14,15]. One of the underlying mechanisms may be the disruption of long and short range projections in the prefrontal and frontal striatal cortex areas associated with DMN [76]. This would reduce the connectivity in the prefrontal cortex, which Pagani and colleagues [76] linked to social communications deficits. Interestingly we found a correspondence with our data: SHANK3 was found to be significantly correlated with decrease clusters in the DMN and Pagani et al [76] report that Shank3B^-/-^ mice have a decreased brain volume in DMN areas. The role of SHANK1 in long-range connectivity is still not very well understood, possibly because mutations in this gene may have less of an impact of synapse morphology, therefore causing a less severe phenotype than fellow SHANK family proteins [20]. Nonetheless, it is still known to alter post-synaptic protein composition, thereby reducing the size of the dendritic spines, weakening basal synaptic transmission [20,77]. We did not find any significant correlations between SHANK2 and the brain networks, even though it is known to be associated with ASD phenotypes [19]. This however, does not mean that it is not implicated with ASD. In fact, our results only imply that its expression in the altered GM regions is not associated to any specific network, that is, its effect might be homogeneous across RSNs.

Unlike the other genes considered, MECP2 encodes for a transcription factor. ASD individuals with mutation in MECP2 have been found to have increased cerebellum volume [78]. It is hypothesized that loss of MECP2 leads to weakening of neuronal connections, therefore inhibiting spontaneous synaptic activity and weakening synaptic plasticity [79] which could explain the correlation found with areas of decreased GM.

Recently Liloia et al. [80] demonstrated that co-alteration of GM areas in ASD patients intersect with structural brain connectivity pathways. Particularly DMN was found to have a crucial role in the GM alterations distribution. Similarly, we found that the DMN was correlated with at least one candidate gene in increase and decrease analysis. It is also interesting to note that Liloia et al. (2021) [80] did not find co-alteration areas to be correlated with genetic co-expression connectivity maps. This may be because that co-alteration is a different phenomenon than simple alteration. Another possible explanation is that they correlated one network to the entirety of the genome as found on AHBA, further supporting our rationale of selecting individual genes.

Several limitations can be individuated in this study. First and foremost, in our study the gene data of Allen Human Brain Atlas only included the left hemisphere of the brain for 4 out of the 6 specimens from which the data was collected. We therefore had to assume there was a symmetry between the two hemispheres. In addition to this, our conclusions are based on Yeo’s extended atlas. Owing to random statistical fluctuation and different grouping of brain areas, the results may change slightly when using a different atlas. Moreover, the data for the meta-analysis come from a variety of articles, which all differ slightly in data collection and methodology.

In this study, we only covered 8 out of the many putative ASD genes, with the intent of conducting an exploratory analysis on some of the genes that might be more representative of the disease. As the results of that, our results clearly are not representative of the whole relation between ASD genetics and GM alterations. However, the rationale behind such approach made it possible to focus our attention on the results about selected genes, instead of dealing with the complexity of big data. The method presented in the study could be used to explore the relationship between a larger set of genes and brain networks to provide further insight into this complex disease. It may also be applied to other disorders with a genetic basis such as schizophrenia.

Having individuated the correlation between specific genes and networks, it would be interesting to investigate how mutations in such genes specifically affect each network both structurally and functionally.

## Supplementary Materials

**Figure S1:**
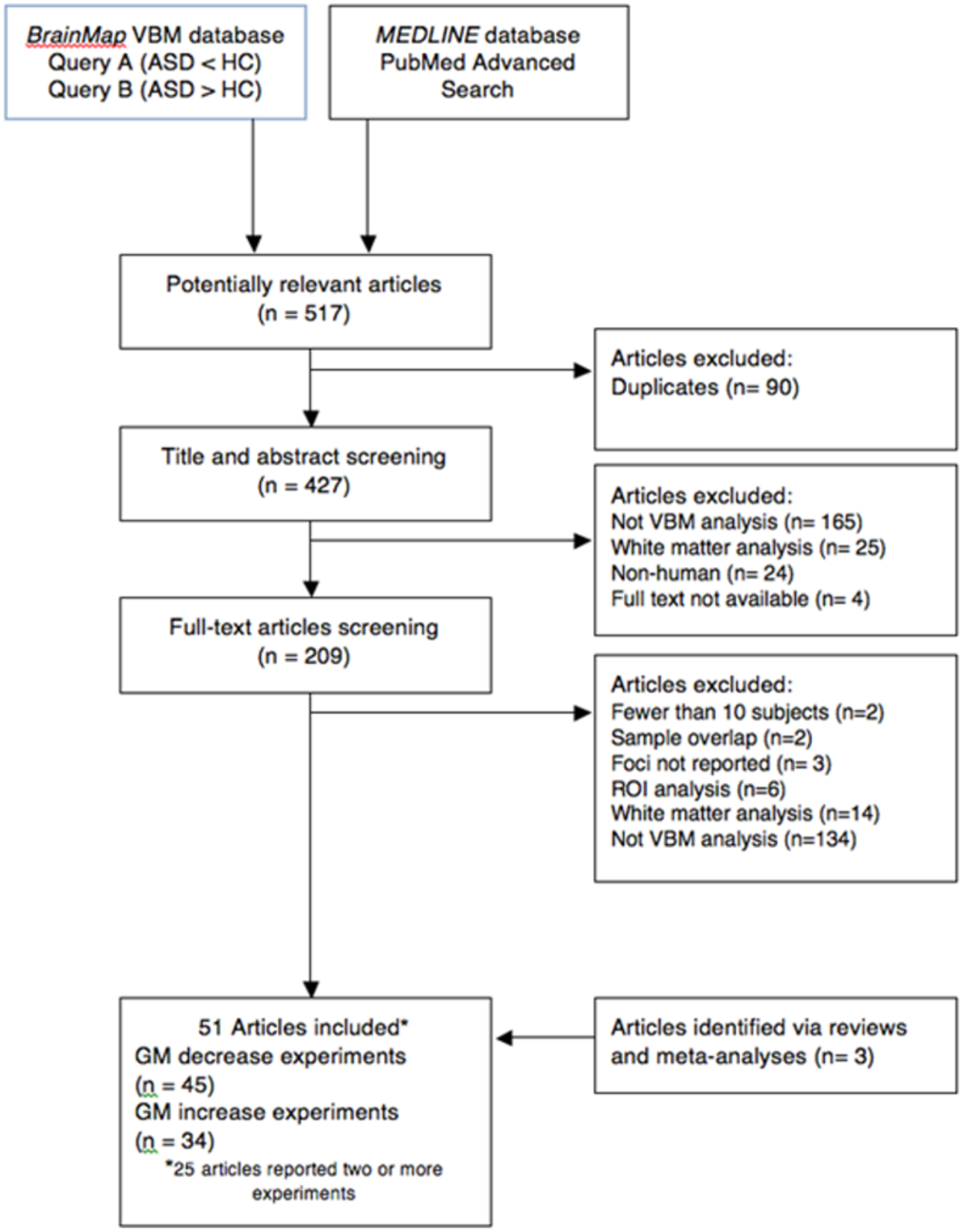
PRISMA flow chart

## Author Contributions

A.C. and E.P. wrote the final draft. T.C, A.C, E.P and L.M designed the study, analyzed the data. D.L, M.F., F.C. and J.M helped in the study design and took part in analysis of the imaging data. All authors have read and agreed to the published version of the manuscript.

## References

1. Baio, J.; Wiggins, L.; Christensen, D.L.; Maenner, M.J.; Daniels, J.; Warren, Z.; Kurzius-Spencer, M.; Zahorodny, W.; Rosenberg, C.R.; White, T.; et al. Prevalence of autism spectrum disorder among children aged 8 Years - Autism and developmental disabilities monitoring network, 11 Sites, United States, 2014. MMWR Surveill. Summ. 2018, 67, doi:10.15585/mmwr.ss6706a1.

2. American Psychiatric Association Diagnostic and Statistical Mental Disorders (Dsm 5). In American Psychiatric Association; 2013.

3. Ansel, A.; Rosenzweig, J.P.; Zisman, P.D.; Melamed, M.; Gesundheit, B. Variation in gene expression in autism spectrum disorders: An extensive review of transcriptomic studies. Front. Neurosci. 2017, 10.

4. Rylaarsdam, L.; Guemez-Gamboa, A. Genetic Causes and Modifiers of Autism Spectrum Disorder. Front. Cell. Neurosci. 2019, 13.

5. Ashburner, J.; Friston, K.J. Voxel-based morphometry - The methods. Neuroimage 2000, 11, doi:10.1006/nimg.2000.0582.

6. Cauda, F.; Geda, E.; Sacco, K.; D’Agata, F.; Duca, S.; Geminiani, G.; Keller, R. Grey matter abnormality in autism spectrum disorder: An activation likelihood estimation meta-analysis study. J. Neurol. Neurosurg. Psychiatry 2011, 82, doi:10.1136/jnnp.2010.239111.

7. Carlisi, C.O.; Norman, L.; Murphy, C.M.; Christakou, A.; Chantiluke, K.; Giampietro, V.; Simmons, A.; Brammer, M.; Murphy, D.G.; Mataix-Cols, D.; et al. Comparison of neural substrates of temporal discounting between youth with autism spectrum disorder and with obsessive-compulsive disorder. Psychol. Med. 2017, 47, doi:10.1017/S0033291717001088.

8. Deramus, T.P.; Kana, R.K. Anatomical likelihood estimation meta-analysis of grey and white matter anomalies in autism spectrum disorders. NeuroImage Clin. 2015, 7, doi:10.1016/j.nicl.2014.11.004.

9. Lukito, S.; Norman, L.; Carlisi, C.; Radua, J.; Hart, H.; Simonoff, E.; Rubia, K. Comparative meta-analyses of brain structural and functional abnormalities during cognitive control in attention-deficit/hyperactivity disorder and autism spectrum disorder. Psychol. Med. 2020, 50.

10. Nickl-Jockschat, T.; Habel, U.; Maria Michel, T.; Manning, J.; Laird, A.R.; Fox, P.T.; Schneider, F.; Eickhoff, S.B. Brain structure anomalies in autism spectrum disorder-a meta-analysis of VBM studies using anatomic likelihood estimation. Hum. Brain Mapp. 2012, 33, doi:10.1002/hbm.21299.

11. Krumm, N.; O’Roak, B.J.; Shendure, J.; Eichler, E.E. A de novo convergence of autism genetics and molecular neuroscience. Trends Neurosci. 2014, 37.

12. Grasby, K.L.; Jahanshad, N.; Painter, J.N.; Colodro-Conde, L.; Bralten, J.; Hibar, D.P.; Lind, P.A.; Pizzagalli, F.; Ching, C.R.K.; McMahon, M.A.B.; et al. The genetic architecture of the human cerebral cortex. Science (80-.). 2020, 367, doi:10.1126/science.aay6690.

13. Lau, H.Y.G.; Fornito, A.; Fulcher, B.D. Scaling of gene transcriptional gradients with brain size across mouse development. Neuroimage 2021, 224, doi:10.1016/j.neuroimage.2020.117395.

14. Guang, S.; Pang, N.; Deng, X.; Yang, L.; He, F.; Wu, L.; Chen, C.; Yin, F.; Peng, J. Synaptopathology involved in autism spectrum disorder. Front. Cell. Neurosci. 2018, 12.

15. He, Y.; Zhou, Y.; Ma, W.; Wang, J. An integrated transcriptomic analysis of autism spectrum disorder. Sci. Rep. 2019, 9, doi:10.1038/s41598-019-48160-x.

16. Bassi, M.S.; Iezzi, E.; Gilio, L.; Centonze, D.; Buttari, F. Synaptic plasticity shapes brain connectivity: Implications for network topology. Int. J. Mol. Sci. 2019, 20.

17. Monteiro, P.; Feng, G. SHANK proteins: Roles at the synapse and in autism spectrum disorder. Nat. Rev. Neurosci. 2017, 18.

18. Durand, C.M.; Betancur, C.; Boeckers, T.M.; Bockmann, J.; Chaste, P.; Fauchereau, F.; Nygren, G.; Rastam, M.; Gillberg, I.C.; Anckarsäter, H.; et al. Mutations in the gene encoding the synaptic scaffolding protein SHANK3 are associated with autism spectrum disorders. Nat. Genet. 2007, 39, doi:10.1038/ng1933.

19. Leblond, C.S.; Heinrich, J.; Delorme, R.; Proepper, C.; Betancur, C.; Huguet, G.; Konyukh, M.; Chaste, P.; Ey, E.; Rastam, M.; et al. Genetic and functional analyses of SHANK2 mutations suggest a multiple hit model of autism spectrum disorders. PLoS Genet. 2012, 8, doi:10.1371/journal.pgen.1002521.

20. Sato, D.; Lionel, A.C.; Leblond, C.S.; Prasad, A.; Pinto, D.; Walker, S.; O’Connor, I.; Russell, C.; Drmic, I.E.; Hamdan, F.F.; et al. SHANK1 deletions in males with autism spectrum disorder. Am. J. Hum. Genet. 2012, 90, doi:10.1016/j.ajhg.2012.03.017.

21. Jamain, S.; Quach, H.; Betancur, C.; Råstam, M.; Colineaux, C.; Gillberg, C.; Soderstrom, H.; Giros, B.; Leboyer, M.; Gillberg, C.; et al. Mutations of the X-linked genes encoding neuroligins NLGN3 and NLGN4 are associated with autism. Nat. Genet. 2003, 34, doi:10.1038/ng1136.

22. Marro, S.G.; Chanda, S.; Yang, N.; Janas, J.A.; Valperga, G.; Trotter, J.; Zhou, B.; Merrill, S.; Yousif, I.; Shelby, H.; et al. Neuroligin-4 Regulates Excitatory Synaptic Transmission in Human Neurons. Neuron 2019, 103, doi:10.1016/j.neuron.2019.05.043.

23. Tabuchi, K.; Blundell, J.; Etherton, M.R.; Hammer, R.E.; Liu, X.; Powell, C.M.; Südhof, T.C. A neuroligin-3 mutation implicated in autism increases inhibitory synaptic transmission in mice. Science (80-.). 2007, 318, doi:10.1126/science.1146221.

24. Canali, G.; Garcia, M.; Hivert, B.; Pinatel, D.; Goullancourt, A.; Oguievetskaia, K.; Saint-Martin, M.; Girault, J.A.; Faivre-Sarrailh, C.; Goutebroze, L. Genetic variants in autism-related CNTNAP2 impair axonal growth of cortical neurons. Hum. Mol. Genet. 2018, 27, doi:10.1093/hmg/ddy102.

25. Gao, R.; Piguel, N.H.; Melendez-Zaidi, A.E.; Martin-de-Saavedra, M.D.; Yoon, S.; Forrest, M.P.; Myczek, K.; Zhang, G.; Russell, T.A.; Csernansky, J.G.; et al. CNTNAP2 stabilizes interneuron dendritic arbors through CASK. Mol. Psychiatry 2018, 23, doi:10.1038/s41380-018-0027-3.

26. Chao, H.T.; Zoghbi, H.Y.; Rosenmund, C. MeCP2 Controls Excitatory Synaptic Strength by Regulating Glutamatergic Synapse Number. Neuron 2007, 56, doi:10.1016/j.neuron.2007.08.018.

27. Na, E.S.; Nelson, E.D.; Kavalali, E.T.; Monteggia, L.M. The impact of MeCP2 loss-or gain-of-function on synaptic plasticity. Neuropsychopharmacology 2013, 38.

28. Raichle, M.E. A paradigm shift in functional brain imaging. J. Neurosci. 2009, 29.

29. Damoiseaux, J.S.; Rombouts, S.A.R.B.; Barkhof, F.; Scheltens, P.; Stam, C.J.; Smith, S.M.; Beckmann, C.F. Consistent restingstate networks across healthy subjects. Proc. Natl. Acad. Sci. U. S. A. 2006, 103, doi:10.1073/pnas.0601417103.

30. Shehzad, Z.; Kelly, A.M.C.; Reiss, P.T.; Gee, D.G.; Gotimer, K.; Uddin, L.Q.; Lee, S.H.; Margulies, D.S.; Roy, A.K.; Biswal, B.B.; et al. The resting brain: Unconstrained yet reliable. Cereb. Cortex 2009, 19, doi:10.1093/cercor/bhn256.

31. Zuo, X.N.; Kelly, C.; Adelstein, J.S.; Klein, D.F.; Castellanos, F.X.; Milham, M.P. Reliable intrinsic connectivity networks: Test-retest evaluation using ICA and dual regression approach. Neuroimage 2010, 49, doi:10.1016/j.neuroimage.2009.10.080.

32. Smith, S.M.; Jenkinson, M.; Woolrich, M.W.; Beckmann, C.F.; Behrens, T.E.J.; Johansen-Berg, H.; Bannister, P.R.; De Luca, M.; Drobnjak, I.; Flitney, D.E.; et al. Advances in functional and structural MR image analysis and implementation as FSL. In Proceedings of the NeuroImage; 2004; Vol. 23.

33. Pievani, M.; Filippini, N.; Van Den Heuvel, M.P.; Cappa, S.F.; Frisoni, G.B. Brain connectivity in neurodegenerative diseases - From phenotype to proteinopathy. Nat. Rev. Neurol. 2014, 10.

34. Cauda, F.; Nani, A.; Manuello, J.; Premi, E.; Palermo, S.; Tatu, K.; Duca, S.; Fox, P.T.; Costa, T. Brain structural alterations are distributed following functional, anatomic and genetic connectivity. Brain 2018, 141, doi:10.1093/brain/awy252.

35. Seeley, W.W.; Crawford, R.K.; Zhou, J.; Miller, B.L.; Greicius, M.D. Neurodegenerative Diseases Target Large-Scale Human Brain Networks. Neuron 2009, 62, doi:10.1016/j.neuron.2009.03.024.

36. Shafiei, G.; Markello, R.D.; Makowski, C.; Talpalaru, A.; Kirschner, M.; Devenyi, G.A.; Guma, E.; Hagmann, P.; Cashman, N.R.; Lepage, M.; et al. Spatial Patterning of Tissue Volume Loss in Schizophrenia Reflects Brain Network Architecture. Biol. Psychiatry 2020, 87, doi:10.1016/j.biopsych.2019.09.031.

37. Sha, Z.; Wager, T.D.; Mechelli, A.; He, Y. Common Dysfunction of Large-Scale Neurocognitive Networks Across Psychiatric Disorders. Biol. Psychiatry 2019, 85, doi:10.1016/j.biopsych.2018.11.011.

38. Hull, J. V.; Jacokes, Z.J.; Torgerson, C.M.; Irimia, A.; Van Horn, J.D.; Aylward, E.; Bernier, R.; Bookheimer, S.; Dapretto, M.; Gaab, N.; et al. Resting-state functional connectivity in autism spectrum disorders: A review. Front. Psychiatry 2017, 7.

39. Sporns, O.; Bassett, D.S. Editorial: New Trends in Connectomics. Netw. Neurosci. 2018, 2.

40. Glahn, D.C.; Winkler, A.M.; Kochunov, P.; Almasy, L.; Duggirala, R.; Carless, M.A.; Curran, J.C.; Olvera, R.L.; Laird, A.R.; Smith, S.M.; et al. Genetic control over the resting brain. Proc. Natl. Acad. Sci. U. S. A. 2010, 107, doi:10.1073/pnas.0909969107.

41. Padmanabhan, A.; Lynch, C.J.; Schaer, M.; Menon, V. The Default Mode Network in Autism. Biol. Psychiatry Cogn. Neurosci. Neuroimaging 2017, 2.

42. Shen, E.H.; Overly, C.C.; Jones, A.R. The Allen Human Brain Atlas. Comprehensive gene expression mapping of the human brain. Trends Neurosci. 2012, 35.

43. Hawrylycz, M.J.; Lein, E.S.; Guillozet-Bongaarts, A.L.; Shen, E.H.; Ng, L.; Miller, J.A.; Van De Lagemaat, L.N.; Smith, K.A.; Ebbert, A.; Riley, Z.L.; et al. An anatomically comprehensive atlas of the adult human brain transcriptome. Nature 2012, 489, doi:10.1038/nature11405.

44. Fortune, S. Voronoi diagrams and delaunay triangulations. In Handbook of Discrete and Computational Geometry, Third Edition; 2017.

45. Torta, D.M.E.; Costa, T.; Duca, S.; Fox, P.T.; Cauda, F. Parcellation of the cingulate cortex at rest and during tasks: A meta-analytic clustering and experimental study. Front. Hum. Neurosci. 2013, doi:10.3389/fnhum.2013.00275.

46. Fox, P.T.; Lancaster, J.L. Mapping context and content: The BrainMap model. Nat. Rev. Neurosci. 2002, doi:10.1038/nrn789.

47. Fox, P.T.; Laird, A.R.; Fox, S.P.; Fox, P.M.; Uecker, A.M.; Crank, M.; Koenig, S.F.; Lancaster, J.L. BrainMap taxonomy of experimental design: Description and evaluation. In Proceedings of the Human Brain Mapping; 2005.

48. Vanasse, T.J.; Fox, P.M.; Barron, D.S.; Robertson, M.; Eickhoff, S.B.; Lancaster, J.L.; Fox, P.T. BrainMap VBM: An environment for structural meta-analysis. Hum. Brain Mapp. 2018, doi:10.1002/hbm.24078.

49. Lancaster, J.L.; Laird, A.R.; Fox, P.M.; Glahn, D.E.; Fox, P.T. Automated analysis of meta-analysis networks. In Proceedings of the Human Brain Mapping; 2005.

50. Liberati, A.; Altman, D.G.; Tetzlaff, J.; Mulrow, C.; Gøtzsche, P.C.; Ioannidis, J.P.A.; Clarke, M.; Devereaux, P.J.; Kleijnen, J.; Moher, D. The PRISMA statement for reporting systematic reviews and meta-analyses of studies that evaluate health care interventions: explanation and elaboration. In Proceedings of the Journal of clinical epidemiology; 2009; Vol. 62.

51. Moher, D.; Liberati, A.; Tetzlaff, J.; Altman, D.G.; Altman, D.; Antes, G.; Atkins, D.; Barbour, V.; Barrowman, N.; Berlin, J.A.; et al. Preferred reporting items for systematic reviews and meta-analyses: The PRISMA statement. PLoS Med. 2009, 6.

52. Müller, V.I.; Cieslik, E.C.; Serbanescu, I.; Laird, A.R.; Fox, P.T.; Eickhoff, S.B. Altered brain activity in unipolar depression revisited: Meta-analyses of neuroimaging studies. JAMA Psychiatry 2017, 74, doi:10.1001/jamapsychiatry.2016.2783.

53. Tahmasian, M.; Sepehry, A.A.; Samea, F.; Khodadadifar, T.; Soltaninejad, Z.; Javaheripour, N.; Khazaie, H.; Zarei, M.; Eickhoff, S.B.; Eickhoff, C.R. Practical recommendations to conduct a neuroimaging meta-analysis for neuropsychiatric disorders. Hum. Brain Mapp. 2019, 40.

54. Laird, A.R.; Robinson, J.L.; McMillan, K.M.; Tordesillas-Gutiérrez, D.; Moran, S.T.; Gonzales, S.M.; Ray, K.L.; Franklin, C.; Glahn, D.C.; Fox, P.T.; et al. Comparison of the disparity between Talairach and MNI coordinates in functional neuroimaging data: Validation of the Lancaster transform. Neuroimage 2010, 51, doi:10.1016/j.neuroimage.2010.02.048.

55. Laird, A.R.; Eickhoff, S.B.; Kurth, F.; Fox, P.M.; Uecker, A.M.; Turner, J.A.; Robinson, J.L.; Lancaster, J.L.; Fox, P.T. ALE meta-analysis workfl ows via the BrainMap database: Progress towards a probabilistic functional brain atlas. Front. Neuroinform. 2009, 3, doi:10.3389/neuro.11.023.2009.

56. Laird, A.R.; Fox, P.M.; Price, C.J.; Glahn, D.C.; Uecker, A.M.; Lancaster, J.L.; Turkeltaub, P.E.; Kochunov, P.; Fox, P.T. ALE meta-analysis: Controlling the false discovery rate and performing statistical contrasts. In Proceedings of the Human Brain Mapping; 2005.

57. Turkeltaub, P.E.; Eden, G.F.; Jones, K.M.; Zeffiro, T.A. Meta-analysis of the functional neuroanatomy of single-word reading: Method and validation. Neuroimage 2002, doi:10.1006/nimg.2002.1131.

58. Eickhoff, S.B.; Laird, A.R.; Grefkes, C.; Wang, L.E.; Zilles, K.; Fox, P.T. Coordinate-based activation likelihood estimation meta-analysis of neuroimaging data: A random-effects approach based on empirical estimates of spatial uncertainty. Hum. Brain Mapp. 2009, doi:10.1002/hbm.20718.

59. Thomas Yeo, B.T.; Krienen, F.M.; Sepulcre, J.; Sabuncu, M.R.; Lashkari, D.; Hollinshead, M.; Roffman, J.L.; Smoller, J.W.; Zöllei, L.; Polimeni, J.R.; et al. The organization of the human cerebral cortex estimated by intrinsic functional connectivity. J. Neurophysiol. 2011, 106, doi:10.1152/jn.00338.2011.

60. Fan, L.; Li, H.; Zhuo, J.; Zhang, Y.; Wang, J.; Chen, L.; Yang, Z.; Chu, C.; Xie, S.; Laird, A.R.; et al. The Human Brainnetome Atlas: A New Brain Atlas Based on Connectional Architecture. Cereb. Cortex 2016, 26, doi:10.1093/cercor/bhw157.

61. Buckner, R.L.; Andrews-Hanna, J.R.; Schacter, D.L. The Brain’s Default Network. Ann. N. Y. Acad. Sci. 2008, 1124, doi:10.1196/annals.1440.011.

62. Washington, S.D.; Gordon, E.M.; Brar, J.; Warburton, S.; Sawyer, A.T.; Wolfe, A.; Mease-Ference, E.R.; Girton, L.; Hailu, A.; Mbwana, J.; et al. Dysmaturation of the default mode network in autism. Hum. Brain Mapp. 2014, 35, doi:10.1002/hbm.22252.

63. Allen, G.; Courchesne, E. Attention function and dysfunction in autism. Front. Biosci. 2001, 6.

64. Farrant, K.; Uddin, L.Q. Atypical developmental of dorsal and ventral attention networks in autism. Dev. Sci. 2016, 19, doi:10.1111/desc.12359.

65. Bednarz, H.M.; Kana, R.K. Patterns of Cerebellar Connectivity with Intrinsic Connectivity Networks in Autism Spectrum Disorders. J. Autism Dev. Disord. 2019, 49, doi:10.1007/s10803-019-04168-w.

66. Bi, X.A.; Zhao, J.; Xu, Q.; Sun, Q.; Wang, Z. Abnormal functional connectivity of resting state network detection based on linear ICA analysis in autism spectrum disorder. Front. Physiol. 2018, 9, doi:10.3389/fphys.2018.00475.

67. Cerliani, L.; Mennes, M.; Thomas, R.M.; Di Martino, A.; Thioux, M.; Keysers, C. Increased functional connectivity between subcortical and cortical resting-state networks in Autism spectrum disorder. JAMA Psychiatry 2015, 72, doi:10.1001/jamapsychiatry.2015.0101.

68. Gutierrez, R.C.; Hung, J.; Zhang, Y.; Kertesz, A.C.; Espina, F.J.; Colicos, M.A. Altered synchrony and connectivity in neuronal networks expressing an autism-related mutation of neuroligin 3. Neuroscience 2009, 162, doi:10.1016/j.neuroscience.2009.04.062.

69. Nguyen, T.A.; Lehr, A.W.; Roche, K.W. Neuroligins and Neurodevelopmental Disorders: X-Linked Genetics. Front. Synaptic Neurosci. 2020, 12.

70. Scott-Van Zeeland, A.A.; Abrahams, B.S.; Alvarez-Retuerto, A.I.; Sonnenblick, L.I.; Rudie, J.D.; Ghahremani, D.; Mumford, J.A.; Poldrack, R.A.; Dapretto, M.; Geschwind, D.H.; et al. Altered functional connectivity in frontal lobe circuits is associated with variation in the autism risk gene CNTNAP. Sci. Transl. Med. 2010, 2, doi:10.1126/scitranslmed.3001344.

71. Ching, M.S.L.; Shen, Y.; Tan, W.H.; Jeste, S.S.; Morrow, E.M.; Chen, X.; Mukaddes, N.M.; Yoo, S.Y.; Hanson, E.; Hundley, R.; et al. Deletions of NRXN1 (neurexin-1) predispose to a wide spectrum of developmental disorders. Am. J. Med. Genet. Part B Neuropsychiatr. Genet. 2010, 153, doi:10.1002/ajmg.b.31063.

72. Barnea-Goraly, N.; Kwon, H.; Menon, V.; Eliez, S.; Lotspeich, L.; Reiss, A.L. White matter structure in autism: Preliminary evidence from diffusion tensor imaging. Biol. Psychiatry 2004, 55, doi:10.1016/j.biopsych.2003.10.022.

73. Frith, C. Is autism a disconnection disorder? Lancet Neurol. 2004, 3.

74. Just, M.A.; Cherkassky, V.L.; Keller, T.A.; Minshew, N.J. Cortical activation and synchronization during sentence comprehension in high-functioning autism: Evidence of underconnectivity. Brain 2004, 127, doi:10.1093/brain/awh199.

75. Geschwind, D.H.; Levitt, P. Autism spectrum disorders: developmental disconnection syndromes. Curr. Opin. Neurobiol. 2007, 17.

76. Pagani, M.; Bertero, A.; Liska, A.; Galbusera, A.; Sabbioni, M.; Barsotti, N.; Colenbier, N.; Marinazzo, D.; Scattoni, M.L.; Pasqualetti, M.; et al. Deletion of autism risk gene shank3 disrupts prefrontal connectivity. J. Neurosci. 2019, 39, doi:10.1523/JNEUROSCI.2529-18.2019.

77. Hung, A.Y.; Futai, K.; Sala, C.; Valtschanoff, J.G.; Ryu, J.; Woodworth, M.A.; Kidd, F.L.; Sung, C.C.; Miyakawa, T.; Bear, M.F.; et al. Smaller dendritic spines, weaker synaptic transmission, but enhanced spatial learning in mice lacking Shank1. J. Neurosci. 2008, 28, doi:10.1523/JNEUROSCI.3032-07.2008.

78. Hashem, S.; Nisar, S.; Bhat, A.A.; Yadav, S.K.; Azeem, M.W.; Bagga, P.; Fakhro, K.; Reddy, R.; Frenneaux, M.P.; Haris, M. Genetics of structural and functional brain changes in autism spectrum disorder. Transl. Psychiatry 2020, 10.

79. Gonzales, M.L.; LaSalle, J.M. The role of MeCP2 in brain development and neurodevelopmental disorders. Curr. Psychiatry Rep. 2010, 12.

80. Liloia, D.; Mancuso, L.; Uddin, L.Q.; Costa, T.; Nani, A.; Keller, R.; Manuello, J.; Duca, S.; Cauda, F. Gray Matter Abnormalities Follow Non-Random Patterns of Co-Alteration in Autism: Meta-Connectomic Evidence. NeuroImage Clin. 2021, doi:10.1016/j.nicl.2021.102583.

